# GpsB is an accessory Z-ring anchor

**DOI:** 10.1101/2024.05.11.593517

**Authors:** Dipanwita Bhattacharya, Lily McKnight, Asher King, Prahathees J. Eswara

## Abstract

Bacterial cytokinesis is a well-coordinated process in which multiple proteins, collectively called the divisome, are recruited to the site of division. A key member of the divisome is FtsZ which forms a ring-like structure (Z-ring) to mark the division site through its association with Z-ring anchors. Synchronized movement of Z-ring filaments and peptidoglycan synthesis along the axis of division help generate a division septum to separate the daughter cells. Thus, FtsZ needs to be linked to the PG synthesis machinery. GpsB is a highly conserved protein among the members of the Firmicutes phylum which has been shown to regulate cell wall synthesis through interaction with penicillin binding proteins. Previously published data from our lab established GpsB as a member of the divisome which directly interacts with FtsZ in *Staphylococcus aureus*. More specifically, we showed that GpsB binds to FtsZ by recognizing the R-X-X-R sequence in its C- terminal tail (CTT) region. As the GpsB recognition sequence is also present in *Bacillus subtilis*, we speculated that GpsB-FtsZ interaction might occur in this organism as well. In support of our prediction, previous studies reported that disruption of *gpsB* and *ezrA* or *gpsB* and *ftsA* is deleterious. Given that both EzrA and FtsA are known Z-ring anchors, we hypothesized that in the absence of other FtsZ anchors, GpsB can fulfill this role in *B. subtilis*. Our data, taken together, conclusively shows that GpsB is a Z-ring anchor in *B. subtilis* and this role of GpsB only becomes apparent in the absence of other FtsZ anchoring proteins. Based on the conserved nature of the R-X-X-R sequence in FtsZ- CTT, we also tested GpsB-FtsZ interaction in *Enterococcus faecalis* and *Listeria monocytogenes* and show that this recognition motif dependent interaction is present in the former but not the latter. Our results suggest the possibility of C-terminal R-X-X-R independent interaction in *L. monocytogenes* and *Streptococcus pneumoniae*. Thus, it appears that GpsB may serve as an accessory Z-ring anchor in multiple organisms.

**Importance:** Cell division is essential for production of offsprings and propagation of life. In bacteria, a key tubulin-like cell division protein FtsZ is brought to the division site by the action of multiple regulators. Arrival of FtsZ and activation of cell wall synthesis are needed to build a division septum to separate the daughter cells. As such, this vital process is controlled by redundant fail-safe mechanisms. For instance, multiple proteins can anchor FtsZ to the membrane at the division site. In this report, we reveal that yet another protein, GpsB, could serve as a back-up FtsZ anchor. GpsB is normally associated with cell wall synthesis in most organisms of the Firmicutes phylum. However, in the absence of other known dedicated FtsZ anchoring factors, GpsB is able to step in to rescue cell division. We provide evidence that this GpsB-FtsZ interaction is present in *Staphylococcus aureus*, *Bacillus subtilis*, *Enterococcus faecalis*, *Listeria monocytogenes*, and possibly *Streptococcus pneumoniae*. As the rapid rise in antibiotic resistance is threatening public health globally, knowledge of unique important cell division factors could be harnessed to develop new antibacterial therapeutics.

## Introduction

Most bacteria divide by a process called binary fission in which the division of parental cell produces two nearly identical daughter cells^1–4^. Although various studies conducted in the past decades elucidated the mechanism of bacterial cell division in great detail, many factors remain elusive^5–7^. One of the most studied cell division proteins is FtsZ, a tubulin-like GTPase that marks the site of cell division^8–10^. FtsZ is conserved in almost all bacteria, archaea^10, 11^, and in eukaryotic organelles mitochondria and chloroplasts^12, 13^. The assembly of FtsZ into FtsZ ring (Z-ring) is spatiotemporally regulated by certain proteins that help tether FtsZ to cell membrane^3^. These Z-ring anchors play a vital role in facilitating the cell division process.

GpsB is a Firmicutes-specific protein that has been shown to interact with a variety of proteins involved in cell wall synthesis^14, 15^. Our group previously showed that *Staphylococcus aureus* GpsB is capable of directly interacting with FtsZ and stimulating its GTPase activity^16^. We also showed that GpsB interacts with FtsZ through a conserved R-X-X-R recognition sequence (S-R-R-T-R-R) located in the C-terminal tail (CTT) region of FtsZ^17^. A positive interaction between FtsZ and GpsB has not been reported in other organisms^14^. However, we observed that R-X-X-R motif is present in the CTT region of other FtsZ homologs in *Bacillus subtilis*, *Enterococcus faecalis*, *Listeria monocytogenes*, but is absent in *Streptococcus pneumoniae* (**Fig. 1A**). Based on this, we speculated that GpsB-FtsZ interaction may be conserved beyond *S. aureus*. The CTT region is a known hot spot for Z-ring anchors such as FtsA, EzrA, and SepF^18^. Thus, we wondered whether GpsB-FtsZ interaction or GpsB-related division phenotype is masked by the presence of other Z-ring anchors. Interestingly, it has been reported that deletion or depletion of either *gpsB* and *ezrA* or *gpsB* and *ftsA* are synthetic sick/lethal in *B. subtilis*^19, 20^. Therefore, it appears that GpsB must be able to compensate for the lack of EzrA or FtsA and serve as a back-up Z-ring anchor.

**Figure 1.**
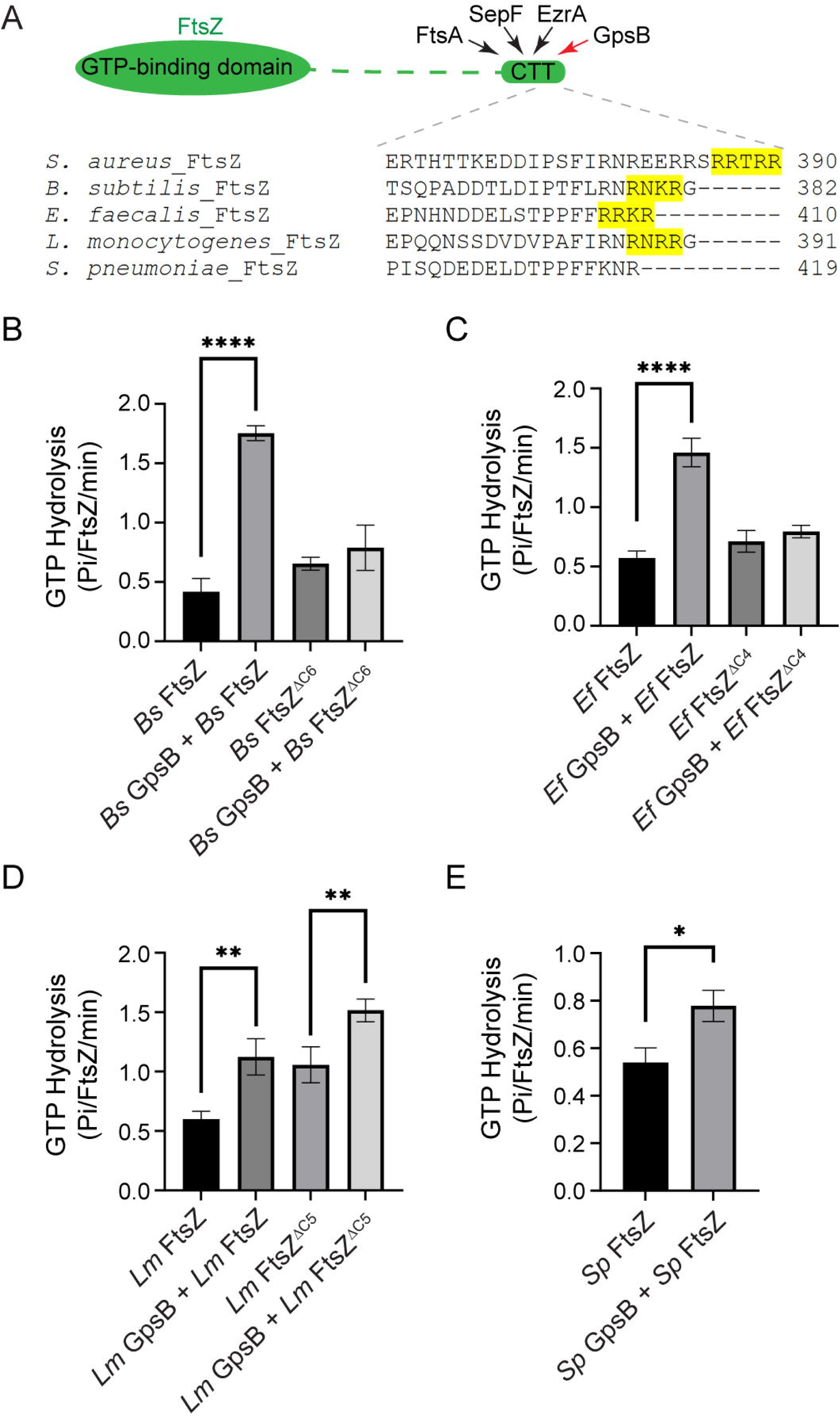
GpsB stimulates the GTPase of FtsZ in multiple species. **(A)** Cartoon representation of FtsZ domains. FtsZ anchoring proteins FtsA, SepF, EzrA, and GpsB (protein of interest in this study) interact with the C-terminal tail of FtsZ. The disordered linker connecting N-terminal domain GTPase domain and C-terminal tail region are shown in dotted lines. Multiple sequence alignment of the C-terminal end of FtsZ from representative species of Firmicutes phylum is provided. The GpsB recognition motif (R-X-X-R) is highlighted. **(B-E)** GTPase activity analyses of full length and C-terminal truncated *Bs* FtsZ (B), *Ef* FtsZ (C), *Lm* FtsZ (D) and *Sp* FtsZ (E) in the absence and presence of GpsB. Error bars indicate standard deviation and n=4. ****, ** and * indicate P<0.0001, 0.001 and 0.01 respectively.

In this study, we sought to investigate GpsB-FtsZ interaction in *B. subtilis* and other species. Biochemically, we were able to show that *B. subtilis* GpsB (*Bs* GpsB) induces the GTPase activity of *B. subtilis* FtsZ (*Bs* FtsZ) in a CTT dependent manner, as showed previously for *S. aureus* GpsB/FtsZ pair^16, 17^. This was the case for *E. faecalis* and *L. monocytogenes* GpsB and FtsZ, however the latter was not dependent on R-X-X-R motif. To our surprise, we also detected a modest but reproducible stimulation of *S. pneumoniae* FtsZ GTPase activity in the presence of GpsB, even though R-X-X-R sequence is absent in this organism. Thus, it is possible another interaction surface may be present to facilitate GpsB-FtsZ interaction in *L. monocytogenes* and *S. pneumoniae*. Other partners of GpsB, such as *B. subtilis* EzrA and MreC^14, 15, 19^, do not have a R-X-X-R motif so presence of this motif is not strictly required for recognition.

It has been shown that deletion of *gpsB* in *B. subtilis* does not exhibit any cell division phenotype under normal growth conditions except in the absence of EzrA or FtsA^19, 20^. We posited that GpsB-dependent cell division phenotypes would become more prominent in the absence of other FtsZ anchors such as EzrA, SepF, or FtsA^3^; as well as in the absence of known interaction partner PBP1^19, 21^. Using fluorescence microscopy, we noticed that increased production of GpsB in otherwise wild type (WT) cells does not lead to cell division inhibition. In contrast, cell elongation was observed in strains lacking EzrA or SepF. Using a temperature sensitive *ftsA* (*ftsA\**) mutant^22^, we show that cell elongation at non-permissive temperature is suppressed by ectopic expression of *gpsB*. Furthermore, we provide evidence that in a *ponA* (which encodes PBP1) deletion background, increased production of GpsB leads to cell division inhibition suggesting PBP1 is an efficient inhibitor of excess GpsB. We also show that GpsB facilitates cell division, as increased cell lengths of Δ*ezrA* Δ*ponA*, Δ*sepF* Δ*ponA*, and *ftsA\** Δ*ponA* are greatly reduced upon overexpression of *gpsB*. Recent reports in *S. aureus* suggest that FtsZ treadmilling kinetics is unaffected in the absence of GpsB^23^, and that GpsB plays a preferential role in peptidoglycan synthesis^23, 24^. Our data show that the GpsB phenotype is more severe in the absence of another FtsZ anchor, EzrA, in *S. aureus* similar to our observation in *B. subtilis*. Therefore, we believe the role of GpsB as a Z-ring anchor is masked by the presence of other FtsZ anchors. Thus, taken together, our data establishes GpsB as an accessory Z-ring anchor in multiple organisms.

## Results

### GpsB stimulates the GTPase activity of FtsZ in multiple species

The CTT of FtsZ is a magnet for its partners. The mechanistic details of interaction between FtsA^18, 25, 26^, EzrA^27, 28^, and SepF^29, 30^ with CTT of FtsZ has been uncovered (**Fig. 1A**). We recently showed that *S. aureus* GpsB also targets CTT of FtsZ for interaction by recognizing the R-X-X-R motif^17^. We also showed that *S. aureus* GpsB stimulates the GTPase activity of FtsZ^16^, in a manner dependent on the terminal R-X-X-R motif^17^. Furthermore, we provided evidence that the binding affinity between GpsB and FtsZ is lost upon truncation of the C-terminal 6 residues in FtsZ^17^. Upon generating a multiple sequence alignment of the CTT residues of different FtsZ homologs of representative bacterial species of Firmicutes phylum, we noticed the presence of R-X-X-R sequence in *B. subtilis*, *E. faecalis*, *L. monocytogenes*, but not in *S. pneumoniae* (**Fig. 1A**).

To test whether GpsB-mediated stimulation of GTPase activity of FtsZ occurs in other species, we purified recombinant GpsB/FtsZ pairs of other organisms. We also tested the truncated FtsZ versions lacking the RXXR sequence. As shown in **Fig. 1B**, co-incubation of *B. subtilis* GpsB (*Bs* GpsB) and *B. subtilis* FtsZ (*Bs* FtsZ) stimulated the GTPase activity by nearly 4-fold. This GpsB-dependent enhancement was absent upon removal of the C-terminal 6 residues in FtsZ (*Bs* FtsZ^ΔC6^), similar to what we reported for *S. aureus* GpsB/FtsZ pair^17^. GpsB alone did not hydrolyze GTP (**Fig. S1A**). Next, we tested *E. faecalis* GpsB/FtsZ proteins (*Ef* GpsB/*Ef* FtsZ). Our results reveal that *Ef* GpsB stimulates the GTPase activity of *Ef* FtsZ approximately 3-fold, but not in the absence of R-X-X-R (**Fig. 1C**). *L. monocytogenes* GpsB (*Lm* GpsB) also stimulated the *L. monocytogenes* FtsZ (*Lm* FtsZ) GTPase activity by nearly 2-fold (**Fig. 1D**). However, this was not R-X-X-R dependent, as truncated FtsZ (*Lm* FtsZ^ΔC5^) by itself displayed higher GTP hydrolysis rate and addition of *Lm* GpsB increased the rate further by ∼1.5-fold. This data suggests that *Lm* GpsB interaction with *Lm* FtsZ is not solely reliant on the R-X-X-R motif. Finally, we tested the GTPase activity stimulation of *S. pneumoniae* FtsZ (*Sp* FtsZ), which naturally lacks R-X-X-R sequence, by *S. pneumoniae* GpsB (*Sp* GpsB). We observed a modest but statistically significant and reproducible increase in GTPase activity in the presence of *Sp* GpsB (**Fig. 1E**). This result indicates that *Sp* FtsZ and *Sp* GpsB may also be partners but do not require R-X-X-R for interaction. Overall, this set of results indicates that the GpsB-FtsZ interaction we initially observed in *S. aureus* is also conserved in *B. subtilis*, *E. faecalis*, *L. monocytogenes*, and *S. pneumoniae*. However, the GpsB-FtsZ interaction in the latter two appears to be independent of R-X-X-R recognition motif.

### Binding of GpsB and FtsZ is widely conserved

Using the purified recombinant GpsB/FtsZ protein pairs, we conducted a fluorescence-based binding assay. Simply, we labeled FtsZ with fluorescein isothiocyanate (FITC) and monitored the change in fluorescence in the presence of equimolar GpsB. It has been previously shown that a change in fluorescence represents binding^31^. First, we monitored the background fluorescence spectra of FITC-labeled *Bs* FtsZ by itself (**Figs. 2A and 2F**). Next, we measured the fluorescence spectra of *Bs* FtsZ mixed with *Bs* GpsB and noted more than double the peak fluorescence intensity. On the contrary, fluorescence spectra of *Bs* FtsZ^ΔC6^ and *Bs* FtsZ^ΔC6^ + *Bs* GpsB did not show any measurable difference in the fluorescence intensity, suggesting R-X-X-R motif in *Bs* FtsZ is needed for binding. This trend is also seen in *Sa* FtsZ/*Sa* GpsB and *Sa* FtsZ^ΔC6^/*Sa* GpsB pairs (**Figs. 2B and 2F**), consistent with what we observed previously^17^. Further, to determine the binding efficiency, a fixed concentration of FITC-*Bs* FtsZ was incubated with increasing concentrations of GpsB. The fluorescence intensity of FITC labeled *Bs* FtsZ increased significantly upon incubating with increasing concentrations of *Bs* GpsB, indicating that *Bs* GpsB specifically interacts with *Bs* FtsZ **(Fig. S2A)**. However, deletion of C-terminal 6 amino acid residues from *Bs* FtsZ abolished its interaction with *Bs* GpsB as no change in the fluorescence intensity of FITC-*Bs* FtsZ^ΔC6^ could be observed upon incubation with *Bs* GpsB **(Fig. S2C)**. Using the change in the fluorescence of FITC-*Bs* FtsZ in the presence of *Bs* GpsB, the K_d_ for the binding of *Bs* GpsB with *Bs* FtsZ was determined to be 48 ± 2 µM **(Fig. S2B)**. After similar analysis of *Sa* FtsZ/*Sa* GpsB pair, we established the K_d_ for their interaction to be 53.81 ± 8 µM (**Fig. S2DE**), which is comparable to the Kd value obtained by surface plasmon resonance previously (40 ± 2 µM)^17^. Again, we failed to see a robust concentration-dependent increase in fluorescence signal between *Sa* FtsZ^ΔC6^ and *Sa* GpsB (**Fig. S2F**). To ensure the specificity of this binding assay, we tested the interaction of FtsZ and an unrelated protein (bovine serum albumin; BSA), which did not result in an increase in fluorescence signal for any of the FtsZ versions tested (**Fig. S2G**).

**Figure 2.**
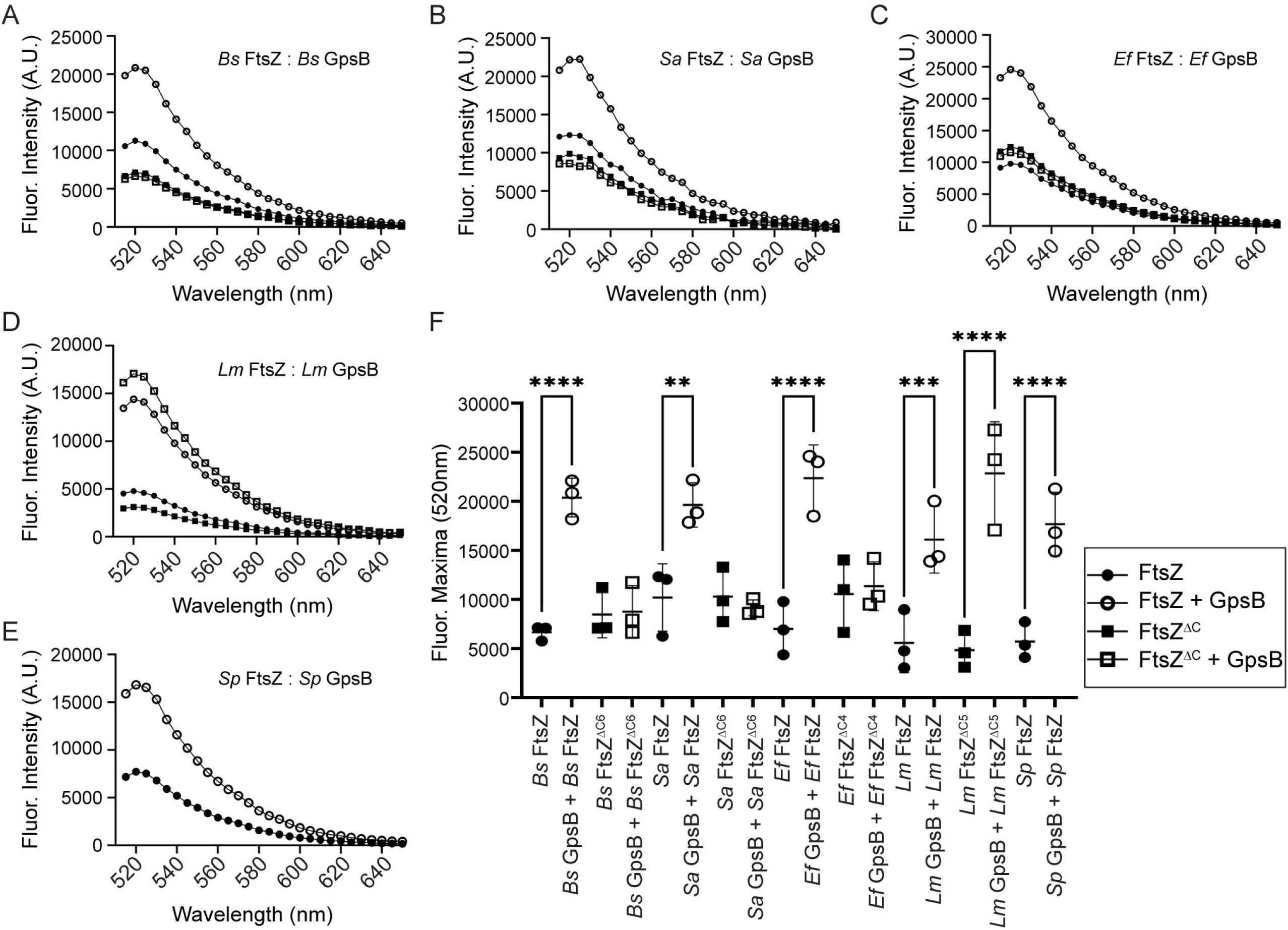
Binding of GpsB and FtsZ is widely conserved. The binding of GpsB to either full length or C-terminal truncated FtsZ of *Bs* **(A),** *Sa* **(B),** *Ef* **(C),** *Lm* **(D)** and *Sp* **(E)** was monitored using fluorescence spectroscopy. The representative fluorescence spectra of full length FITC-FtsZ (1 µM) in the absence (●) and presence (○) of GpsB (1 µM) were plotted against the wavelength. Similarly, truncated version of FITC-FtsZ (1 µM) was incubated without (▪) or with (□) GpsB and the fluorescence spectra were monitored. (**F**) The average of maximum fluorescence intensities at 520 nm; n=4 and **, *** and **** indicate p ∼ 0.0025, 0.0006 and <0.0001 respectively.

We also investigated the interaction between the FtsZ/GpsB pairs of other species. We noticed a significant increase in fluorescence intensity between FITC-*Ef* FtsZ and *Ef* GpsB to a similar magnitude observed for *B. subtilis* and *S. aureus* proteins (**Figs. 2C and 2F**). This enhancement in fluorescence intensity was absent when *Ef* GpsB was incubated with FITC-*Ef* FtsZ^ΔC4^. This result is consistent with the GTPase assay results (**Fig. 1C**). This suggests that the interaction between *Ef* FtsZ and *Ef* GpsB is mediated by the 4 C-terminal amino acids of *Ef* FtsZ. Next, we monitored the fluorescence intensity of FITC-*Lm* FtsZ and FITC-*Lm* FtsZ^ΔC5^ with or without *Lm* GpsB (**Figs. 2D and 2F**). We observed an increase in fluorescence intensity for FITC-*Lm* FtsZ and *Lm* GpsB consistent with other pairs. However, in striking contrast, we also noticed an increase in fluorescence intensity for FITC-*Lm* FtsZ^ΔC5^ and *Lm* GpsB. We noted previously that stimulation of GTPase activity was also seen when *Lm* FtsZ^ΔC5^ was incubated with *Lm* GpsB (**Fig. 1D**), which is not the case for other species. This set of data reveals that in *L. monocytogenes*, the interaction between FtsZ and GpsB is conserved but independent of the signature GpsB recognition (R-X-X-R) motif located within the 5 C-terminal residues of *Lm* FtsZ. Finally, we investigated the fluorescence spectra of *Sp* GpsB incubated with FITC-*Sp* FtsZ, which lacks the R-X-X-R motif in the C-terminus, and noted statistically significant increase in peak fluorescence intensity (**Fig. 2EF**). This is consistent with the stimulation of GTPase activity of *Sp* FtsZ by *Sp* GpsB (**Fig. 1E**).

Taken together, the binding of GpsB to FtsZ through the R-X-X-R motif was confirmed in *B. subtilis* and *E. faecalis* using fluorescence assay. However, GpsB-FtsZ interaction in *L. monocytogenes* and *S. pneumoniae*, although present, appears to be independent of R-X-X-R motif. Thus, the interaction between GpsB and FtsZ that we first observed in *S. aureus*^17^ is also present in multiple species.

### Effects of GpsB are only apparent in the absence of known FtsZ anchors

It is known that under standard growth conditions, there are no discernible *gpsB* phenotypes in *B. subtilis*^19, 20^. Based on our results confirming that GpsB interacts with FtsZ, we speculated that potential *gpsB* phenotypes are masked by the presence of redundant FtsZ anchors such as SepF, FtsA, and EzrA that all target the same binding site – CTT region on FtsZ (**Fig. 1A**). Therefore, we hypothesized that *gpsB* phenotypes will become more apparent in the absence of other FtsZ targeting proteins. To test this, we constructed a *B. subtilis* strain harboring an inducible copy of *gpsB* at an ectopic locus and investigated the *gpsB* overexpression phenotype in Δ*sepF* and Δ*ezrA* strains. We also tested this in a temperature sensitive *ftsA* mutant (*ftsA*\*; FtsA^S9N^)^22^ background grown at non-permissive temperature, as deletion of *ftsA* in *B. subtilis* leads to severe/lethal filamentation^30, 32, 33^. Overexpression of *gpsB* by itself did not have any noticeable effect on cell length (**Fig. 3AB**; compare the cell lengths in the absence and presence of inducer). Although Δ*sepF* or Δ*ezrA* cells are longer in general, overexpression of *gpsB* in these strain backgrounds lead to further increase in cell length suggesting that excess GpsB is negatively affecting FtsZ ring assembly. Similar cell elongation was observed when *ezrA* or *sepF* is overexpressed^34–36^. In contrast, we see that the ectopic expression of *gpsB* rescues cell division when *ftsA*\* cells are grown at non-permissive temperature which suggests that GpsB can step in as a Z-anchor when FtsA is nonfunctional (**Fig. S3AB**). We also tested whether GpsB overproduction would have any effect on cells lacking ZapA, which does not interact with the C-terminal tail region of FtsZ^35, 37^, and thus not compete with GpsB. As shown in **Fig. S3C**, overexpression of *gpsB* in Δ*zapA* strain background resulted in modest but statistically significant cell length increase. In addition, we investigated the *gpsB* phenotypes in Δ*facZ* background, as FacZ also has a role in proper Z-ring assembly^38^. Although Δ*facZ* cells are longer as previously reported, our results reveal that deletion of *gpsB* or overexpression of *gpsB* does not affect cell length further (**Fig. S3D**). This data underscores a possible accessory role of GpsB in *B. subtilis* cell division in the absence of the other known FtsZ anchors.

**Figure 3.**
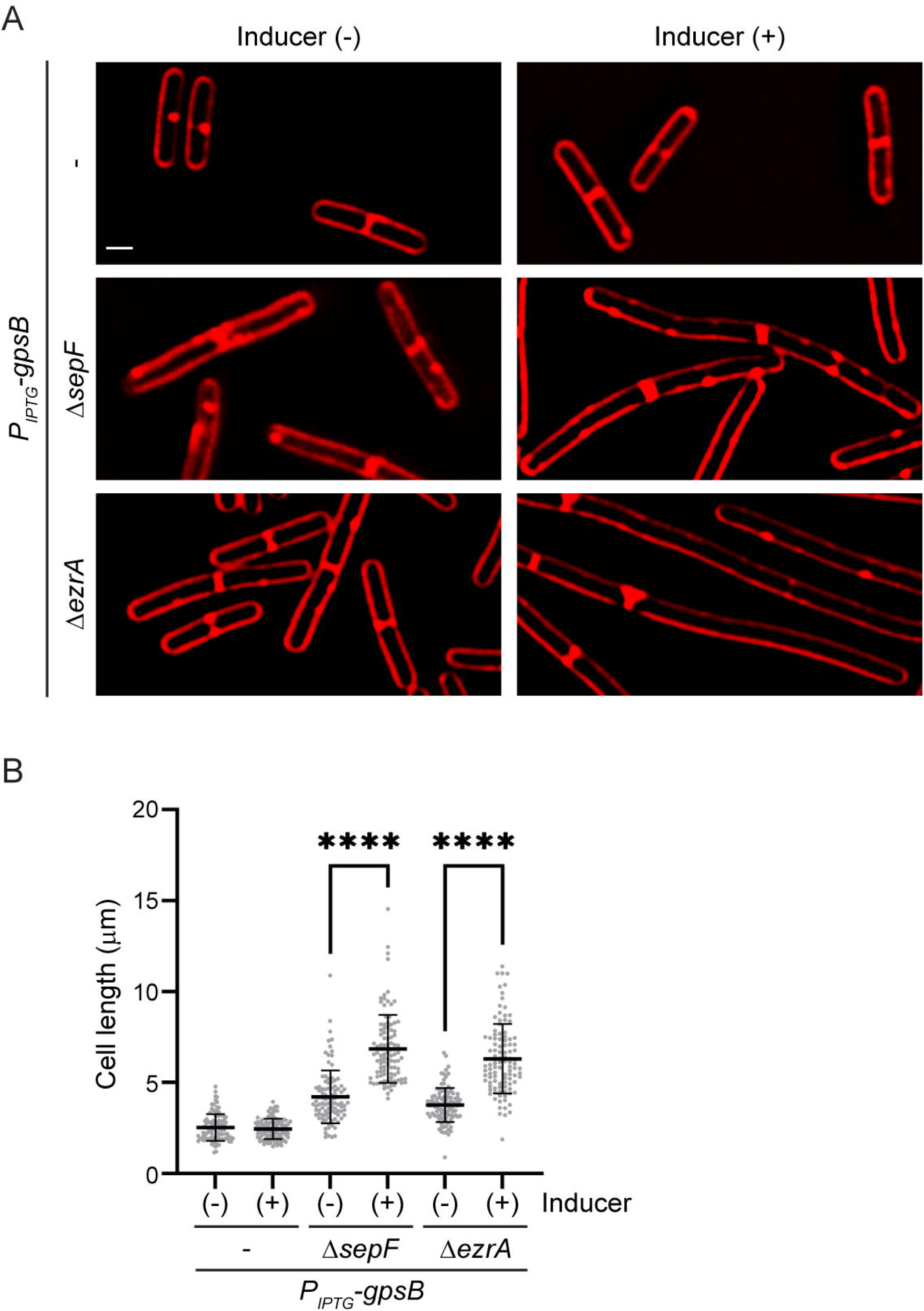
GpsB overproduction worsens the cell elongation phenotypes of strains lacking SepF or EzrA. **(A)** The effect of *gpsB* overexpression in *B. subtilis* in otherwise WT (GG18), Δ*sepF* (BDB12), and Δ*ezrA* (BDB1) backgrounds. Micrographs of cells grown in the absence (left) or presence (right) of the inducer are shown. Cell membrane was stained with synapto-red. Scale bar is 1 µm. (B) Quantification of the cell lengths of *B. subtilis* strains shown in panel A. **** indicates p< 0.0001; n=100.

### GpsB serves as an accessory FtsZ anchor in the absence of its partner PBP1

GpsB is known to interact with penicillin binding proteins (PBPs) in *B. subtilis*, *S. aureus*, and other organisms by recognizing R-X-X-R motif^14, 15, 17, 21^. Therefore, we hypothesized that in the absence of *ponA* (which encodes GpsB interaction partner, PBP1), GpsB will be more likely to interact with FtsZ as both PBP1 and FtsZ interact with GpsB through their R-X-X-R motifs. In affirmation of our speculation, overexpression of *gpsB* in Δ*ponA* background leads to cell length increase (**Fig. 4AB**), which is not the case in the absence of inducer. Next, we wanted to probe whether GpsB can serve as a FtsZ anchor in the absence of PBP1. To test this, we investigated the effects of *gpsB* overexpression in the absence of PBP1 and one of the known FtsZ anchors. In the absence of *gpsB* induction, deletion of *ponA* and either *sepF* or *ezrA* lead to increased cell lengths. Interestingly, in the presence of inducer, the cell lengths of Δ*ponA* Δ*sepF* and Δ*ponA* Δ*ezrA* were greatly reduced (**Fig. 4AB**). Similarly, cell division inhibition in Δ*ponA ftsA*\* grown at non-permissive temperature was also abrogated by overexpression of *gpsB* (**Fig. S4AB**). Overall, these results suggest that GpsB can in fact facilitate cell division by serving as FtsZ anchor in the absence of other partners of GpsB or FtsZ.

**Figure 4.**
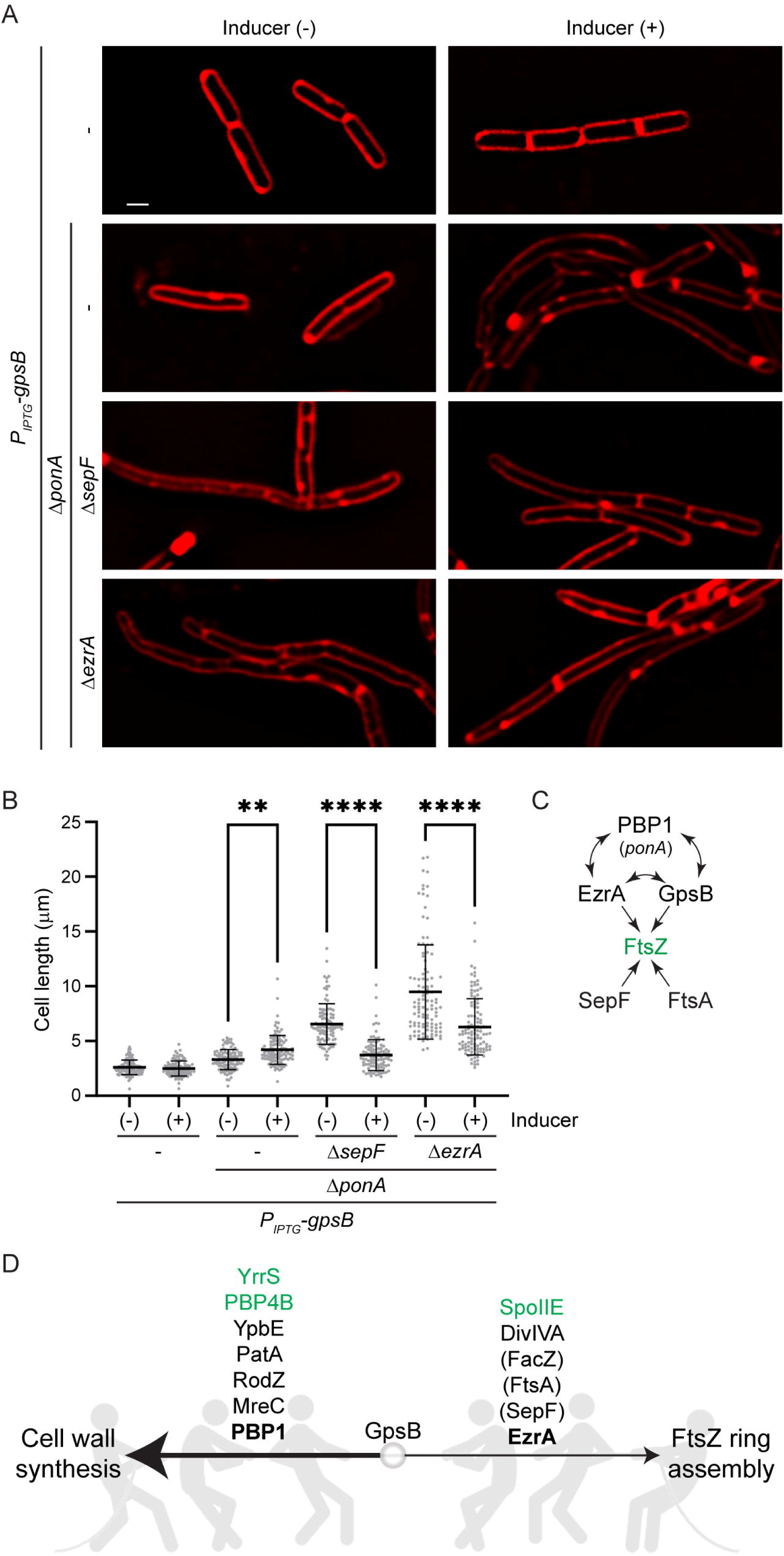
GpsB overproduction phenotypes in Δ*ponA* and upon additional deletion of Δ*sepF* or Δ*ezrA*. **(A)** Micrographs of *gpsB* overexpression strain (GG18), and *gpsB* overexpression in Δ*ponA* (BDB20), Δ*ponA* Δ*sepF* (BDB18), or Δ*ponA* Δ*ezrA* (BDB22) strain backgrounds. Cells were grown in the absence (left) or presence (right) of the inducer. Cell membrane was stained with synapto-red. Scale bar is 1µm. **(B)** Cell length quantification of strains shown in panel A. ** and **** indicate p<0.001 and p<0.0001 respectively; n=100. **(C)** Simplified interaction network of GpsB and Z-anchors. **(D)** Tug-of-war model of GpsB roles. A well-characterized role of GpsB is in regulating cell wall synthesis (thick arrow) which is executed through partners such as PBP1, MreC, and RodZ. The function of GpsB in facilitating Z-ring assembly is minor due to the presence of redundant FtsZ anchors and binding site competition for the FtsZ C-terminal tail. However, the ability of GpsB to anchor FtsZ becomes apparent in the absence of one of the major Z-ring anchors or PBP1. Other GpsB binding partners that may be involved in these processes are also listed. Proteins that are not determined to be direct partners are listed within parentheses. The proteins shown in green are involved in sporulation.

Based on our results, we wondered whether GpsB facilitated cell division in *ftsA*\*, Δ*sepF*, and Δ*ponA* strains. To test this, we deleted *gpsB* in these strain backgrounds. We noticed that the cells lengths of Δ*sepF* Δ*gpsB* and Δ*ponA* Δ*gpsB* were significantly longer than single deletions (**Fig. S4C**). This suggested that indeed GpsB is capable of promoting cell division in these backgrounds. However, we did not notice any difference in cell lengths between *ftsA*\* and *ftsA*\* Δ*gpsB* strains. As *ftsA* and *gpsB* has been reported to be synthetically essential^20^, we suspect that in *ftsA*\* although non-functional FtsA is still present, and thus could occupy and occlude the binding site needed for GpsB to help anchor FtsZ. We were unable to generate a Δ*ezrA* Δ*gpsB* strain without an inducible copy of *gpsB*, consistent with previously reported synthetic sick/lethal phenotype^19^. Therefore, we analyzed the cell lengths of this strain in the absence or presence of inducer. In the absence of inducer, the culture grew poorly, and the cells were long (**Fig. S4D**). However, in the presence of inducer, cell length decreased. Again, this result confirmed that in cells lacking *ezrA*, presence of GpsB is critically needed to facilitate cell division. Overall, our results reveal that GpsB is capable of mediating efficient cell division in the absence of other FtsZ anchors or the well characterized interaction partner of GpsB, PBP1^21^.

### *S. aureus* cells lacking *ezrA* display severe GpsB-dependent cell division phenotype

It has been shown that the division site localization of GpsB depends on EzrA in *S. aureus*^39^. Evidence of direct interaction between GpsB and septum-localized peptidoglycan (PG) synthesis machinery proteins such as PBP2 and PBP4 are also available^17, 23, 40^. Recent reports suggest that the predominant role of GpsB is in maintaining normal *S. aureus* cell morphology by spatially restricting PBP2 and PBP4 enzymes at division site for septal PG synthesis^23, 24^. Therefore, we speculated that in the absence of EzrA, excess GpsB will lead to increased peripheral PG synthesis. As predicted, we see that cells overexpressing *gpsB* in a Δ*ezrA* strain background were significantly enlarged in comparison to the cells overexpressing *gpsB* in an otherwise WT background (**Fig. 5AB**). We infer our results to mean that in the absence of EzrA, GpsB is unable to remain at division site and promote septum-specific PG synthesis. Thus, EzrA is a spatial regulator of GpsB. Recently, Bartlett et al. showed that *ezrA* and a new cell division gene *facZ* are synthetically essential, but *ezrA* and *gpsB* are not^38^. They further showed that Δ*facZ* cells are enlarged only in the presence of GpsB and that FacZ is needed for proper localization of GpsB. Thus, it appears that both EzrA and FacZ are redundant GpsB positioning factors. Based on our results and recent reports from other groups^23, 24, 38^, it appears that the major role of GpsB in *S. aureus* is to restrict PBP2 and PBP4 mediated PG synthesis to division septa. As our data clearly shows *Sa* FtsZ-*Sa* GpsB interaction (**Figs. 1 and 2**)^16, 17^, it is possible that this accessory role becomes important in the absence of other FtsZ anchors such as FtsA/SepF and/or PG synthesis enzymes PBP2/PBP4 or under non-standard growth/stress conditions. However, this remains to be tested.

**Figure 5.**
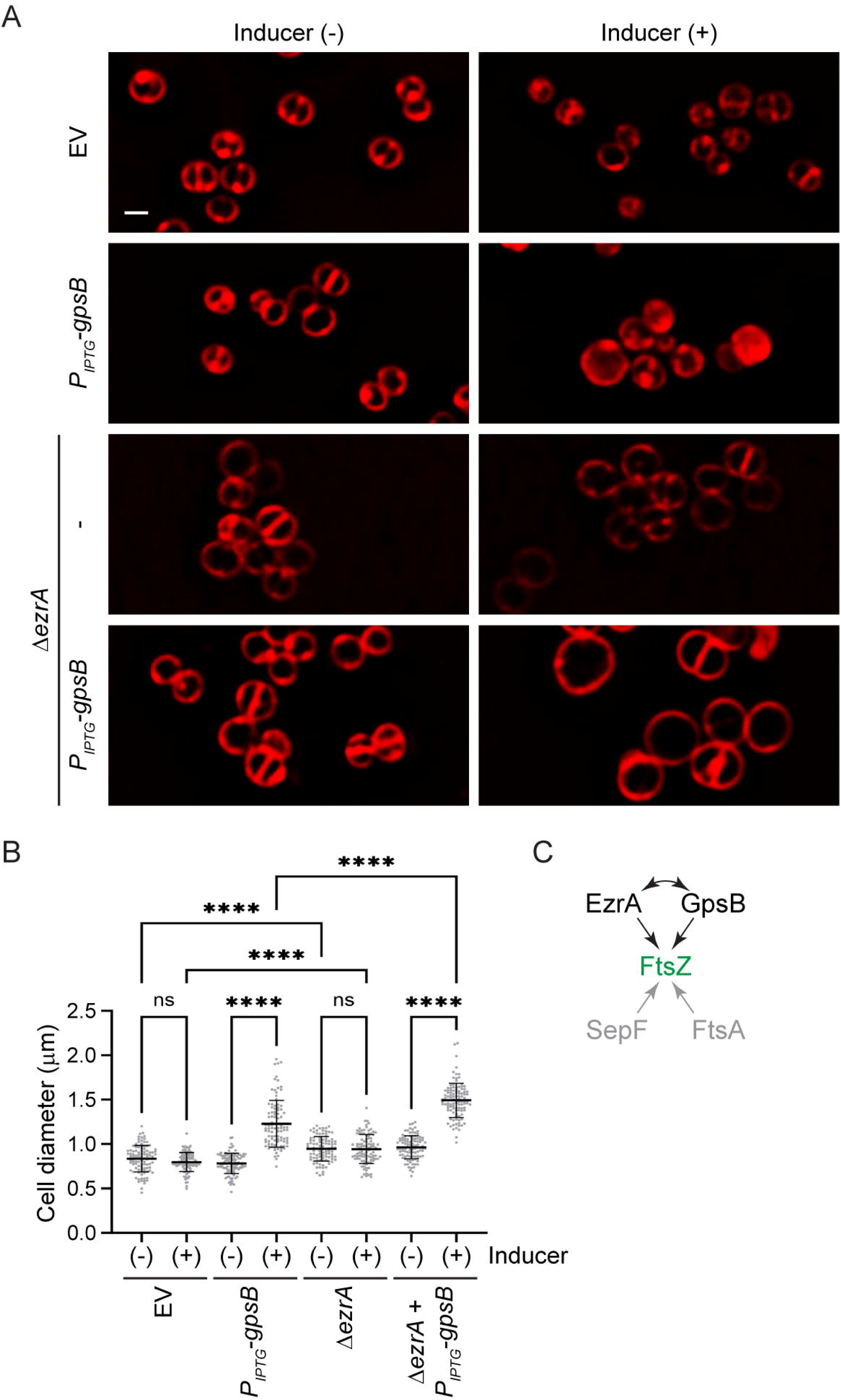
GpsB overexpression in *S. aureus* cells lacking *ezrA* leads to increased cell diameter typical of cell division inhibition. **(A)** The micrographs of *S. aureus* WT cells harboring empty vector (EV; PE355), vector with IPTG-inducible *gpsB* (PE356) and Δ*ezrA* cells harboring EV (LM121) or inducible *gpsB* (LM122) are shown. The cells were grown in the absence (left) and presence (right) of the inducer. **(B)** Cell diameter quantification of the strains shown in panel A. **** indicates p<0.0001; n=100. **(C)** Simplified interaction network of Z-anchors in *S. aureus*.

## Discussion

In this report, we provide evidence showing that GpsB-FtsZ pairs of *B. subtilis* and *E. faecalis* require the presence of C-terminal R-X-X-R recognition sequence in FtsZ for interaction and stimulation of GTPase activity, which is in line with what we previously reported for *S. aureus* GpsB-FtsZ interaction^17^. In contrast, we find that the interaction between GpsB and FtsZ is independent of the terminal R-X-X-R motif in *L. monocytogenes*. This is the case for *S. pneumoniae* which naturally lacks the terminal GpsB recognition motif. However, we cannot rule out the possibility that interaction is mediated by a motif similar to R-X-X-R located in other parts of FtsZ. Overall, our data suggests that GpsB-FtsZ interaction is likely broadly conserved in Firmicutes.

To further validate the physiological relevance of our in vitro results, we focused on FtsZ and GpsB in *B. subtilis*. In this bacterium, it was previously reported that simultaneous deletion/depletion of *gpsB* with either *ezrA* or *ftsA* is synthetically sick/lethal^19, 20^. In addition, enrichment of GpsB at division sites has been noted previously^19, 20, 41^. We find that although overexpression of *gpsB* by itself does not reveal any noticeable phenotype, we see increased cell lengths (cell division inhibition) when *gpsB* is overexpressed in the absence of a functional copy of known FtsZ anchors SepF or EzrA. In contrast, we see that GpsB facilitated division in cells lacking functional FtsA. These results suggested that robust GpsB/FtsZ interaction likely happens only in the absence of other FtsZ tethering proteins due to competition for the same binding site. We also observed that overproduction of GpsB in cells lacking its other interaction partner, PBP1 (which competes with FtsZ using its own R-X-X-R motif^21^), resulted in cell elongation. This suggests that PBP1 is capable of sequestering excess GpsB away from FtsZ. Upon inspecting the cells lacking *ponA* and one of the known FtsZ anchors, we noticed that these cells are longer in the absence of inducer. However, upon GpsB overproduction cell division was restored, and cell elongation was significantly reduced. This data suggested that GpsB is capable of serving as FtsZ anchor in the absence of its partner PBP1 and any one of the known FtsZ anchoring proteins. It has been shown that SepF, a peripheral membrane protein, can anchor FtsZ and cells lacking SepF and either FtsA or EzrA are nonviable^30, 42^. Overproduction of SepF rescues the cell division inhibition seen in cells lacking FtsA^43^. Thus, it is conceivable that in the absence of other factors perhaps GpsB also aids in anchoring FtsZ, as overexpression of *gpsB* restores cell division in the absence of functional FtsA. In support of this notion, we find that cells lacking *ezrA*, *sepF*, or *ponA* are longer and absence of GpsB in these strain backgrounds lead to severe cell division inhibition in the case of Δ*ezrA* and mild but significant cell elongation in the latter two. Therefore, we believe that GpsB is an accessory but bona fide FtsZ anchor.

PBP1 plays a major role in peripheral PG synthesis, as deleting *ponA* restores some of the abnormal cell width phenotypes^44–46^. Besides PBP1 and EzrA, other interaction partners of GpsB include MreC, RodZ, YpbE, and PatA(DapX)^19, 21, 47^. While the first two have clear roles in peripheral cell wall synthesis^48^, the functions of the latter two are less clear. After FtsZ protofilaments arrive at the potential division site, the ring-like structure is further condensed by FtsZ bundling proteins to initiate septal PG synthesis^36, 49^. We identified that another divisome protein, FtsL, is also critical for FtsZ condensation^50^. FtsL controls the functionality of main septal PG synthase machinery made of FtsW and PBP2B (transglycosylase and transpeptidase pair)^50–52^. It was shown that effects of *ezrA* deletion can be countered by *ftsL* overexpression^53^. In fact, FtsL is less stable in cells lacking EzrA^54^. It is known that EzrA and GpsB directly interact with bifunctional PBP1^19, 21, 55^. PBP1 is also a part of the division machinery and is enriched at division sites^56, 57^. It appears that *B. subtilis* cells grown in different media conditions may require PBP1 for proper cell division^58^. It is also to be noted that cell envelopes of Gram-positive monoderm organisms such as *B. subtilis* are composed two distinct layers of PG^59–61^. Thus, cell division in *B. subtilis* utilizes both FtsW/PBP2B as well as PBP1 for efficient cell division. Interestingly, in another member of the Firmicutes phylum *Clostridioides difficile* bifunctional class A PBP (PBP1) is the main driver of septal PG synthesis during vegetative cell division^62^. In *C. difficile*, which does not encode *gpsB*, Z-ring anchors must therefore connect FtsZ to class A PBP1. Bacterial species of other phyla, such as *E. coli* and *Acinetobacter baumannii*, also utilize class A PBP for efficient cell division^63, 64^.

It has been reported that PBP1 localizes to asymmetric cell division sites during sporulation and is required for efficient septation^56^. Evidence showing that GpsB localizes to asymmetric division sites and interacts with another sporulation-specific FtsZ interaction partner, SpoIIE, exists^65^. SpoIIE also interacts with PBP1 and EzrA among others. Thus, it is possible GpsB helps anchor FtsZ for efficient asymmetric cell division during sporulation. It has been reported that GpsB also interacts with YrrS and PBP4B (*yrrR*/*pbpI*) which are produced during sporulation^21^. During vegetative growth, a paralog of GpsB, DivIVA, that controls an FtsZ positioning (Min) system in *B. subtilis*^2, 66^, is also an interaction partner of GpsB^67^. A newly identified factor FacZ, which is found to interact with GpsB in *S. aureus*, appears to also play a role in Z-ring positioning in *B. subtilis*^38^. Taking all of this into account, we propose a tug-of-war model where the role of GpsB in promoting cell wall synthesis and FtsZ ring assembly is influenced by its interaction partners or other factors that compete with GpsB for binding its substrate proteins (**Fig. 4D**).

Previously, we showed that overproduction of *S. aureus* GpsB in *B. subtilis* is lethal due to cell division inhibition^16, 17^. Furthermore, spontaneous suppressor mutations that tolerated lethal *Sa* GpsB overproduction mapped to *tagG* or *tagH* genes that encode wall teichoic acids exporter complex^41^. Here we show that *Sa* GpsB also utilizes R-X-X-R of *B. subtilis* FtsZ for recognition (**Fig. S1**). We believe that the major source of *Sa* GpsB toxicity is due to direct interaction with *Bs* FtsZ, that TagG/TagH mutants circumvent by an unknown mechanism. We suspect that we did not isolate any suppressors harboring mutations in *Bs* FtsZ as the interaction surface is vital for other FtsZ interaction partners.

Recent reports indicate the *S. aureus* GpsB plays a predominant role in spatiotemporal regulation of peptidoglycan synthesis^23, 24^, and that there is no effect on the kinetics of FtsZ ring assembly and constriction^23^. As EzrA^39^ or FacZ^38^ is needed for proper GpsB localization, it could be argued that EzrA and FacZ play a redundant role in restricting GpsB to division sites. Additionally, other regulatory interaction partners of EzrA such as CozEa and CozEb are also present in *S. aureus*^68^. The penicillin binding proteins, PBP2 and PBP4 are interaction partners of GpsB^17, 23^, and are normally enriched at sites of cell division^40^. However, it appears that in the absence of GpsB, PBP2 and PBP4 are no longer enriched at division sites, and they synthesize PG throughout the cell periphery instead^23^. Conversely, overproduction of GpsB leads to increased cell diameter^16^. In this report, we find that this phenotype is exacerbated in the absence of EzrA. Using super-resolution microscopy, we previously showed that GpsB localizes to the leading edge of invaginating membrane during cell division^16^, almost identical to what would be expected of FtsZ during Z-ring constriction. We also uncovered that GpsB is highly capable of directly interacting with FtsZ and stimulating its GTPase activity^16, 17^. Thus in *S. aureus*, GpsB is capable of both (i) stimulating FtsZ ring remodeling and (ii) promoting septal peptidoglycan synthesis needed for the construction of division septum. However, due to redundant well-established Z-ring anchors such as EzrA, SepF, FtsA, and newly uncovered cell division factors such as FacZ^38^ and PcdA^69^, the roles of GpsB in cell division is eclipsed at least under standard laboratory conditions.

Similar to its homolog in other organisms, *L. monocytogenes* GpsB is also targeted to division sites^70^. GpsB of this species interacts with FtsZ anchors SepF and EzrA, as well as several other divisome proteins^21^. *Lm* GpsB is uniquely essential when grown at higher temperature (42 °C), and suppressors have been mapped to pathways that enhance the PG precursor pool^71^. As GpsB interacts with class A PBP A1^21, 70^, it is speculated that in the absence of GpsB, PBP A1 consumes PG precursors for PG synthesis and that increase in PG precursors therefore restores cell division^71^. However, as PBP A1 is required for septal cell wall synthesis^71^, it can be hypothesized that in the absence of GpsB at 42 °C, PBP A1 is dysregulated and functions outside of divisome complex. Therefore, perhaps GpsB directly (**Figs. 1 and 2**) and/or indirectly via interaction with SepF and EzrA coordinates Z-ring remodeling with the action of PBP A1.

The role of GpsB in septal peptidoglycan synthesis has also been established in *S. pneumoniae*^72, 73^. It was shown that GpsB localizes to division sites, and lack of GpsB leads to increased cell length due to aberrant Z-ring assembly^72^. It was proposed that the main role of GpsB is to facilitate septal PG synthesis and restrict peripheral PG synthesis of the elongasome machinery^72, 73^. Thus, it would not be surprising if GpsB was able to interact with FtsZ to execute its role in enabling septal PG synthesis. However, although GpsB and FtsZ are found to be in a complex via co-immunoprecipitation assay^73^, the interaction was absent when tested by bacterial two-hybrid analysis^72, 73^. In this organism, GpsB partner and Z-ring anchor EzrA is essential^74^. Nevertheless, given EzrA localization depends on GpsB^72^, it is possible GpsB-FtsZ interaction may be needed for efficient recruitment of EzrA to the division site and facilitate septal PG synthesis. FtsA is also essential in *S. pneumoniae*, and the aberrantly long cell morphology of cells lacking SepF or GpsB can be rescued by increased levels of FtsA^75^. Therefore, it is conceivable there may be a role for GpsB, albeit minor, in anchoring FtsZ if our biochemical data shown in **Figs. 1 and 2** were to hold true when tested by other methods.

Although the precise function of enterococcal GpsB remains to be understood, it has been shown that cells lacking *gpsB* display cell separation defect and are highly susceptible to cell wall targeting antibiotics^76^. Interestingly, deletion of ser/thr kinase IreK also leads to increased sensitivity to antibiotics, and it was shown that phosphoablative mutant of GpsB is unable to complement Δ*gpsB* phenotype. Thus, it appears the post-translational regulation of GpsB is important for antibiotic resistance. In fact, GpsB of other species are phosphorylated and/or GpsB stimulates the kinase activity of the corresponding ser/thr kinase^14, 15, 76–81^. Thus, it would be interesting to study whether the binding of GpsB and its interaction partners is affected by phosphorylation.

In summary, we show that the interaction between FtsZ and GpsB is conserved in other organisms and may not strictly require R-X-X-R sequence for recognition. As cell division is an essential process, multiple redundant mechanisms are in place, and our data reveals that GpsB also serves as a Z-anchor. However, its significance is only apparent in the absence of one of the other FtsZ anchors or PBP1. The importance of FtsZ-GpsB interaction in *E. faecalis*, *L. monocytogenes*, *S. pneumoniae*, and possibly several other organisms remain to be elucidated.

## MATERIALS AND METHODS

### Strain construction

All *B. subtilis* strains used in this study are derivatives of PY79^82^. The strain details are provided in **Table S1**. To generate *B. subtilis* strains BDB1, BDB12, BDB13, plasmid pGG5^16^ harboring *gpsB* under IPTG-inducible promoter (*gpsB^+^*) was used to clone into the *amyE* locus of the parent strains GG17, RB73, and MW393 lacking *ezrA, sepF*, and harboring temperature sensitive *ftsA^S9N^* respectively. The recombination was confirmed via *amyE* screening. To generate strains lacking *ponA* in the above-mentioned strain backgrounds, the genomic DNA was isolated from strain PE719 and was transformed into BDB1, BDB12 and BDB13 resulting in strains BDB22, BDB18 and BDB24 respectively. In addition, *ΔponA gpsB^+^* strain was constructed by transforming GG18 with the chromosomal DNA from PE482 resulting in strain BDB20. The genomic DNA of GG13 was used to transform into GG18 resulting in strain BDB38. To generate *ΔzapA* strain, PY79 strain was transformed with the chromosomal DNA of RL2647^37^ to generate strain BDB53. The resultant strain was transformed with plasmid pGG5 to construct BDB55. In order to generate double deletion strains, chromosomal DNA of GG13 was used to transform strains RB73 and MW393 resulting in strains BDB31 and BDB34 respectively. As *ΔponA* strain is non-competent, we used PE482 chromosomal DNA to transform GG13 to generate BDB41. We were unable to generate *ΔezrA ΔgpsB* strain without the inducible copy of *gpsB* present, to generate BDB36 we used the chromosomal DNA of GG17 to transform BDB38. To create *ΔfacZ* strains, chromosomal DNA of BKE29780 (Bacillus Genetic Stock Center) was transformed into PY79, GG13 and GG18 to create strains LM138, LM139 and LM140, respectively.

For recombinant protein expression and purification, pET28a plasmid was used to clone the *ftsZ* and *gpsB* genes. The chromosomal DNA from PY79, *L. monocytogenes* EGDe (ATCC strain), *E. faecalis* 29212 (ATCC strain) and *S. pneumoniae* D39 (Chromosomal DNA; gift – Nicholas De Lay lab) were used as templates. The amplified products were ligated to pET28a vector between NdeI and XhoI restriction sites for *ftsZ* and XbaI and BamHI sites for *gpsB*. The final cloned plasmids (from DH5a) were transformed into BL21-DE3 cells for recombinant protein purification. The strains generated and the oligonucleotides used in this study are provided in **Table S1 and S2**. Purified FtsZ and GpsB proteins cloned from different organisms mentioned were used for the biochemical experiments performed in this study.

### Media and growth condition for microscopy

*B. subtilis* cultures were grown overnight at 30 °C in LB medium. Then the cultures were diluted to 1:10 in fresh LB and allowed to grow at 37 °C until mid-log phase (OD_600_ ∼ 0.6-0.8) following which, the cultures were standardized to OD_600_ ∼ 0.1 in fresh LB medium. IPTG was used at 1 mM final concentration at OD_600_ ∼ 0.1 to induce the expression of genes under IPTG-inducible promoter. The cultures were grown for an additional 2 h at 37 °C.

### Recombinant protein purification

BL21-DE3 overexpressing recombinant full length and truncated *ftsZ* were grown until the OD_600_ reached 0.8. Then the cells were induced using IPTG (1 mM for *ftsZ* constructs and 0.5 mM for *gpsB* constructs) and were further allowed to grow at 30 °C for an additional 6 h. Subsequently, the cells were harvested by centrifuging the culture at 8000 rpm for 10 min. The cell pellet was dissolved, homogenized and lysed in ice cold lysis buffer A (20 mM HEPES, pH 7.4; 50 mM KCl; 5 mM MgCl_2_; and 10% glycerol) containing 1 mM PMSF and 1 mg/ml lysozyme. The cell suspension was sonicated, and the lysate was cleared by centrifugation at 35000 rpm for 1 h in an ultracentrifuge. Imidazole (10 mM) was added to the supernatant, and it was loaded to the Ni-NTA column. The column was washed extensively with lysis buffer containing 25 mM and subsequently 50 mM imidazole to elute loosely bound proteins. Both FtsZ and truncated FtsZ were then eluted with lysis buffer containing 250 mM imidazole. The eluted fractions were analyzed via SDS-PAGE and those containing FtsZ were pooled and loaded onto a gravity column containing Sephadex G-25 fine resin (Cytiva) for salt removal. Proteins were eluted in storage buffer containing (20 mM HEPES and 10% glycerol, pH 7.4). FtsZ constructs were then concentrated using Amicon 30 kDa MWCO (Sigma) centrifugal filter and stored at -80°C.

BL21-DE3 cells overexpressing *gpsB* were allowed to grow until OD_600_ 0.6 and then induced with 0.5 mM IPTG and was grown for further 6 h at 37 °C. The cells were harvested, and the protein was purified as described in ^83^.

### GTPase assay

The effect of GpsB on the inorganic phosphate released by FtsZ and truncated FtsZ was examined using malachite green phosphate assay kit (Sigma). Briefly, either FtsZ or truncated FtsZ (30 µM) was incubated with GpsB (10 µM) in 25 mM HEPES buffer containing 140 mM KCl, 5 mM MgCl_2_ and 2 mM GTP at 37 °C for 15 min. Then, the reaction mixtures were incubated with the working reagent provided with the kit for further 30 min in the dark and the absorbance of the reaction milieu were taken at 650 nm^16, 83^.

### Fluorescence microscopy

Fluorescence microscopy was performed as described previously^84^. Briefly, 500 µl aliquots of *B. subtilis* cultures to be imaged were pelleted and washed with 1X PBS. The pellets were resuspended in 100 µl PBS and 1 µg/ml Synapto-Red was added to the cell suspension to stain the cell membrane. Small volume (5 µl) of the suspension was spotted on the glass bottom dish (MatTek) and 1% agarose pad was gently placed on top. The images were taken using DeltaVision Elite deconvolution fluorescence microscope with Photometrics CoolSnap HQ2 camera. Images were deconvoluted by SoftWorx imaging software provided by the manufacturer. The cell lengths were analyzed by ImageJ software and the statistical analysis was carried out using GraphPad Prism.

### Labeling of FtsZ

Both full length and truncated versions of FtsZ (50 µM) was incubated with FITC (250 µM) in 50 mM phosphate buffer (pH 8.0) for 3 h on ice. The reaction was terminated by adding 5 mM Tris-HCl (pH 8.0). The free FITC was separated from the labelled protein by passing the reaction mixture through Sephadex G25 fine column (Cytiva). The concentration of FITC bound FtsZ was determined by taking the absorbance at 495 nm. The concentration of the unlabeled protein was measured using Bradford reagent^85^. The stoichiometry of labelling of FITC per FtsZ was determined to be ∼ 0.7^83^.

### Fluorescence spectroscopy

To check the specificity of binding of GpsB to FtsZ, either FITC-FtsZ (100 nM) or FITC-FtsZ^ΔC6^ (100 nM) was incubated with increasing concentrations of GpsB (10 - 80 µM) in 25 mM HEPES (pH 6.7) for 10 min at 25 °C. The fluorescence spectra of FITC-FtsZ (510-650 nm) were monitored by exciting the reaction mixtures at 495 nm. The changes in the fluorescence intensities of FITC-FtsZ in the presence of different concentrations of GpsB were plotted. The dissociation constants of binding were determined by fitting the values in the following quadratic binding equation^83^.

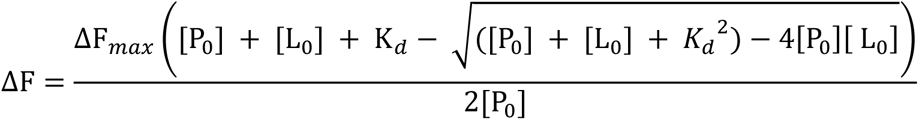

Where ΔF and ΔF_max_ are the change and maximum change in the fluorescence intensity of FITC-FtsZ upon binding to GpsB respectively. P_0_ and L_0_ are the concentrations of FITC-FtsZ and GpsB used, respectively. ΔF was calculated by subtracting the fluorescence intensity of FITC-FtsZ in presence of GpsB from the intensity of the same in the absence of GpsB. The data were analyzed using Graph Pad Prism software.

