## Supplemental Information for "GpsB is an accessory Z-ring anchor"


### Supplemental Table S1

**Strains used in this study**

### Supplemental Table S2

**Oligonucleotides used in this study**

### Supplementary Figure S1

**Cross-species interaction of GpsB and FtsZ. (A)** Both *Sa* GpsB and *Bs* GpsB were found to increase the GTPase activity of *Bs* FtsZ and *Sa* FtsZ respectively. However, GpsB alone individually did not show any GTP hydrolysis. **** indicates P value<0.0001; n=4. **(B)** Cross-species interaction between *Sa* GpsB and *Bs* FtsZ was confirmed by fluorescence spectroscopy. The fluorescence spectra of FITC-*Bs* FtsZ without (●) and with (○) *Sa* GpsB are shown. Similarly, FITC-*Bs*FtsZ$^{\Delta C6}$ was incubated without (■) and with (□) *Sa* GpsB and the fluorescence spectra were graphed. Representative data of four independent spectra recorded is shown.

### Supplementary Figure S2

**GpsB/FtsZ dissociation constant estimation. (A-F)** Both full length (A and D) and truncated (C and F) FITC-labeled *Bs* and *Sa* FtsZ were titrated with a range of GpsB concentrations and the change in the fluorescence (ΔF) were plotted. FITC-FtsZ (100 nM) was incubated without (●) and with 10 μM (■), 15 μM (▲), 20 μM(▼), 40 μM (♦), 50 μM (○), 60 μM (□) and 80 μM (Δ) GpsB and the fluorescence spectra were plotted. (B) and (E) The changes in the fluorescence intensities of FITC labeled full length *Bs* and *Sa* FtsZ in the absence and presence of GpsB were plotted against various concentrations of GpsB used. The dissociation constant of *Bs* FtsZ/GpsB and *Sa* FtsZ/GpsB binding were determined to be 48 ± 2 μM and 54 ± 8 μM respectively. One of the four independent spectra for *B. subtilis* and *S. aureus* are shown. (**G**) The maximum fluorescence intensities of FITC labeled full length FtsZ (1 μM) when incubated without (●) and with (○) bovine serum albumin (BSA) and C-terminal truncated FtsZ without (■) and with (□) BSA (1 μM) at 520 nm were plotted. FtsZ from *B. subtilis*, *S. aureus, E. faecalis, L. monocytogenes*, *and S. pneumoniae* were used and BSA was used as a control.

### Supplementary Figure S3

**GpsB overproduction in temperature-sensitive *ftsA (ftsA*)*, Δ*zapA*, and Δ*facZ* *backgrounds*. (A)** Micrographs of WT (PY79), *ftsA** (MW393), inducible *gpsB* (GG18) and inducible *gpsB* in *ftsA** (BDB13) strains were grown at non-permissive 45 °C without or with inducer. Cell membrane was stained with synapto-red. Scale bar, 1 μm. **(B)** Cell length quantification of the strains shown in panel A. **** indicates p value < 0.0001; n=100. **(C)** Cell lengths of WT (PY79), Δ*zapA* (BDB53), Δ*zapA* with inducible *gpsB* (BDB55) strains. **** indicates p value < 0.0001; n=100. **(D)** Quantification of cell lengths of WT (PY79), Δ*gpsB* (GG13), Δ*facZ* (LM138), Δ*gpsB* Δ*facZ* (LM139), inducible *gpsB* (GG18), and Δ*facZ* with inducible *gpsB* (LM140) strains; n=100. For (C) and (D), cells were grown at 37 °C without or with inducer as indicated in the figure.

### Supplementary Figure S4

**GpsB serves as an accessory Z-ring anchor. (A)** The strains with inducible *gpsB* construct in WT (GG18), Δ*ponA* (BDB20), or *ftsA** Δ*ponA* (BDB24) backgrounds were grown at non-permissive 45 °C without and with inducer. Cell membrane was stained with synapto-red. Scale bar, 1 μm. **(B)** The cell lengths of each strain shown in panel A were quantified. ** and **** indicate p ~ 0.005 and p< 0.0001 respectively; n=100. **(C)** The cell lengths of WT (PY79) and mutants of *ΔgpsB* (GG13), *ftsA** (MW393), *ΔgpsB ftsA** (BDB34), *ΔsepF* (RB73), *ΔgpsB ΔsepF* (BDB31), *ΔponA* (PE719), and *ΔgpsB ΔponA* (BDB41) were quantified and plotted. n=100 and **** indicates p< 0.0001. **(D)** Strains with inducible *gpsB* construct lacking either *gpsB* (BDB38), *ezrA* (BDB1), or both (BDB36) were grown in the absence and presence of IPTG. Note: *ΔgpsB ΔezrA* cells grew very poorly in the absence of inducer. **** indicates p<0.0001; n=100.

**Supplemental Table S1**

| Species | Strain | Genotype | Reference |
|---|---|---|---|
| *B. subtilis* | | | |
| | PY79 | Wild type | [1] |
| | GG13 | *ΔgpsB::tet* | [2] |
| | GG17 | *ΔezrA::spc::cat* | Derived from FG381 (RL2647)[3] |
| | GG18 | *amyE::P*$_{hyperspank}$*-gpsB*$^{BS}$ *spc* | [2] |
| | RB73 | *ΔsepF*::erm | Derived from BGSC, BKE15390 |
| | MW393 | *ftsA*$^{ts}$ *cat* | 1A787 (BGSC); FtsA($^{S9N}$) [4, 5] |
| | PE719 | *ΔponA::kan* | Derived from BGSC, BKK22320 |
| | PE482 | *ΔponA::erm* | Derived from BGSC, BKE22320 |
| | BDB1 | *ΔezrA::spc::cat; amyE::P*$_{hyperspank}$*-gpsB*$^{BS}$ *spc* | This study |
| | BDB12 | *ΔsepF::erm; amyE::P*$_{hyperspank}$*-gpsB*$^{BS}$ *spc* | This study |
| | BDB13 | *ftsA*$^{ts}$ *cat*; *amyE::P*$_{hyperspank}$*-gpsB*$^{BS}$ *spc* | This study |
| | BDB18 | *ΔponA::kan*; *ΔsepF::erm; amyE::P*$_{hyperspank}$*-gpsB*$^{BS}$ *spc* | This study |
| | BDB20 | *ΔponA::erm*; *amyE::P*$_{hyperspank}$*-gpsB*$^{BS}$ *spc* | This study |
| | BDB22 | *ΔponA::kan*; *ΔezrA::spc::cat*; *amyE::P*$_{hyperspank}$*-gpsB*$^{BS}$ *spc* | This study |
| | BDB24 | *ΔponA::kan*; *ftsA*$^{ts}$ *cat*; *amyE::P*$_{hyperspank}$*-gpsB*$^{BS}$ *spc* | This study |
| | BDB31 | *ΔsepF::erm*; *ΔgpsB::tet* | This study |
| | BDB34 | *ftsA*$^{ts}$ *cat*; *ΔgpsB::tet* | This study |
| | BDB36 | *ΔezrA::spc::cat*; *amyE::P*$_{hyperspank}$*-gpsB*$^{BS}$ *spc; ΔgpsB::tet* | This study |
| | BDB38 | *amyE::P*$_{hyperspank}$*-gpsB*$^{BS}$ *spc*; *ΔgpsB::tet* | This study |
| | BDB41 | *ΔponA::erm*; *ΔgpsB::tet* | This study |
| | BDB53 | *ΔzapA-yshB::tet* | Derived from RL2647[3] |
| | BDB55 | *ΔzapA-yshB::tet*; *amyE::P*$_{hyperspank}$*-gpsB*$^{BS}$ *spc* | This study |
| | LM138 | *ΔfacZ::erm* | Derived from BGSC, BKE29780 |
| | LM139 | *ΔfacZ::erm*; *ΔgpsB::tet* | This study |
| | LM140 | *ΔfacZ::erm*; *amyE::P*$_{hyperspank}$*-gpsB*$^{BS}$ *spc* | This study |
| *S. aureus* | | | |
| | PE355 | *S. aureus* RN4220 *harboring pCL15* | [2, 6] |
| | PE356 | *S. aureus* RN4220 *harboring pPE45 (gpsB*$^{SA}$*)* | [2] |
| | LM121 | *S. aureus* RN4220 *ΔezrA harboring pCL15* | This study (Gift - Wenqi Yu lab) |
| | LM122 | *S. aureus* RN4220 *ΔezrA harboring pPE45 (gpsB*$^{SA}$*)* | This study |
| *E. coli* | | | |
| | PE401 | BL21-DE3 harboring pET28a *P*$_{IPTG}$*-gpsBa*$^{Sa}$*-his* | [2] |
| | PE630 | BL21-DE3 harboring pET28a *P*$_{IPTG}$*-his-ftsZ*$^{Sa}$ | [2] |
| | EDB01 | BL21-DE3 harbouring pET28a *P*$_{IPTG}$*-his-ftsZ*$^{Sa(\Delta C6)}$ | [7] |
| | EDB20 | BL21-DE3 harboring pET28a *P*$_{IPTG}$*-his-ftsZ*$^{Bs}$ | This study |
| | EDB27 | Rosetta-DE3 harboring pET28a *P*$_{IPTG}$*-his-ftsZ*$^{Bs(\Delta C6)}$ | This study |
| | EDB24 | Rosetta-DE3 harboring pET28a *P*$_{IPTG}$*-his-gpsB*$^{Bs}$ | This study |
| | EDB33 | BL21-DE3 harboring pET28a P$_{IPTG}$*-his-FtsZ*$^{Lm}$ | This study |
| | EDB35 | BL21-DE3 harboring pET28a *P*$_{IPTG}$*-his-*FtsZ$^{Lm(\Delta C5)}$ | This study |
| | EDB32 | BL21-DE3 harboring pET28a *P*$_{IPTG}$*-his-FtsZ*$^{Ef}$ | This study |

| | | | |
|---|---|---|---|
| | EDB29 | BL21-DE3 harboring pET28a $P_{IPTG}$-*his-FtsZ*$^{Ef(\Delta C4)}$ | This study |
| | EDB34 | BL21-DE3 harboring pET28a $P_{IPTG}$-*his-FtsZ*$^{Sp}$ | This study |
| | EDB30 | BL21-DE3 harboring pET28a $P_{IPTG}$-*his-gpsB*$^{Lm}$ | This study |
| | EDB40 | BL21-DE3 harboring pET28a $P_{IPTG}$-*his-gpsB*$^{Ef}$ | This study |
| | EDB31 | BL21-DE3 harboring pET28a $P_{IPTG}$-*his-gpsB*$^{Sp}$ | This study |

**Supplemental Table S2**

| Primer | Sequence (5’ to 3’) |
|---|---|
| oDB61(*Bs*FtsZ) | AATAA CATATG ATGTTGGAGTTCGAAACAAACATAGACGGCTTAG |
| oDB62(*Bs*FtsZ) | AATAA CTCGAG TTAGCCGCGTTTATTACGGTTTCTTAAGAATGTC |
| oDB68(*Bs*GpsB) | CCC TCTAGA AATAATTTTGTTTAACTTTAAGAAGGAGATATACC ATGCTTGCTG ATAAAGTAAA GCTTTCTGCG AAAGAAATTT |
| oDB69(*Bs*GpsB) | AAA GGATCC TTA ATGATGATGATGATGATG ATCATAAAGCTTGCTGCCAAAAACGTGTTTTTCT |
| oDB34(*Bs*FtsZΔC6) | AATAA CTCGAG TTA TCTTAAGAATGTCGGGATGTCAAGCGTATCATCAGCCGGC |
| oDB119(*Ef*FtsZ) | AATAA CATATG ATGGAATTTTCATTAGACAATAACATTAACAACGGTGCAGT |
| oDB120(*Ef*FtsZ) | AATAA CTCGAG TTATCGTTTTCTGCGGAAAAATGGTGGCGTACTTAATTC |
| oDB121(*Ef*FtsZΔC4) | AATAA CTCGAG TTA GAAAAATGGTGGCGTACTTAATTCGTCATCATTGTGA |
| oDB122(*Ef*GpsB) | CCC TCTAGA_AATAATTTTGTTTAACTTTAAGAAGGAGATATACC ATGGCAAATTTAGTATATAGTCCTAAAGAC |
| oDB123(*Ef*GpsB) | AAA GGATCC TTA ATGATGATGATGATGATG AAATTGACGTGTTTGTGCATTATCAAC |
| oDB124(*Lm*FtsZ) | AATAA CATATG ATGTTAGAATTTGACACTAGTTCAGAAAGTTTGGCAACAAT |
| oDB125(*Lm*FtsZ) | AATAA CTCGAG TTATCCGCGACGGTTACGGTTACGGATAAATGCTGGTAC |
| oDB126(*Lm*FtsZΔC6) | AATAA CTCGAG TTAGTTACGGATAAATGCTGGTACATCAACATCTGAAC |
| oDB127(*Lm*GpsB) | CCC TCTAGA_AATAATTTTGTTTAACTTTAAGAAGGAGATATACC ATGACTTCGGAACAATTTGAGTATCACTTAACAGGCAAAG |
| oDB128(*Lm*GpsB) | AAA GGATCC TTA ATGATGATGATGATGATG TTCGTTATCGTCCAGCTTATTTCCAAAAACATGTTTTTC |
| oDB129(*Sp*GpsB) | CCC TCTAGA_AATAATTTTGTTTAACTTTAAGAAGGAGATATACC ATGGCAAGTATTATTTTTTCAGCGAAAGATATTTTTGAACAAG |
| oDB130(*Sp*GpsB) | AAA GGATCC TTA ATGATGATGATGATGATG AAAATCTGAGTTATCTAAAATTTGTTTACCA |
| oDB88(*Sp*FtsZ) | AATAA CATATG ATGACATTTTCATTTGATACAGCTGCTGCTCAAGGGGC |
| oDB89(*Sp*FtsZ) | AATAA CTCGAG TTAACGATTTTTGAAAAATGGAGGTGTATCCAATTCATCTTCATCTTGTGAAATTGGGGC |

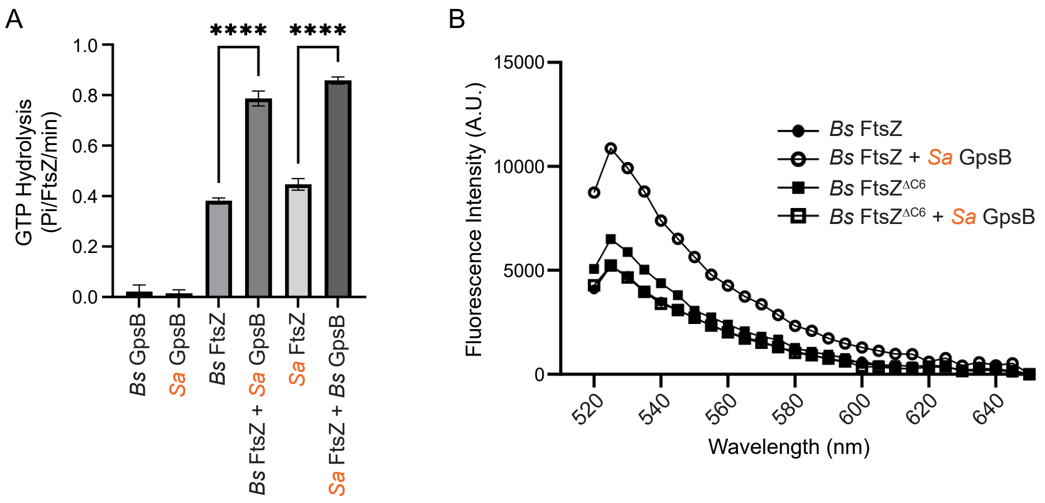

A
GTP Hydrolysis
(Pi/FtsZ/min)
1.0
0.8
0.6
0.4
0.2
0.0
****
****
Bs GpsB
Sa GpsB
Bs FtsZ
Bs FtsZ + Sa GpsB
Sa FtsZ
Sa FtsZ + Bs GpsB
B
Fluorescence Intensity (A.U.)
15000
10000
5000
0
520
540
560
580
600
620
640
Wavelength (nm)
Bs FtsZ
Bs FtsZ + Sa GpsB
Bs FtsZ^ΔC6
Bs FtsZ^ΔC6 + Sa GpsB


Figure S1

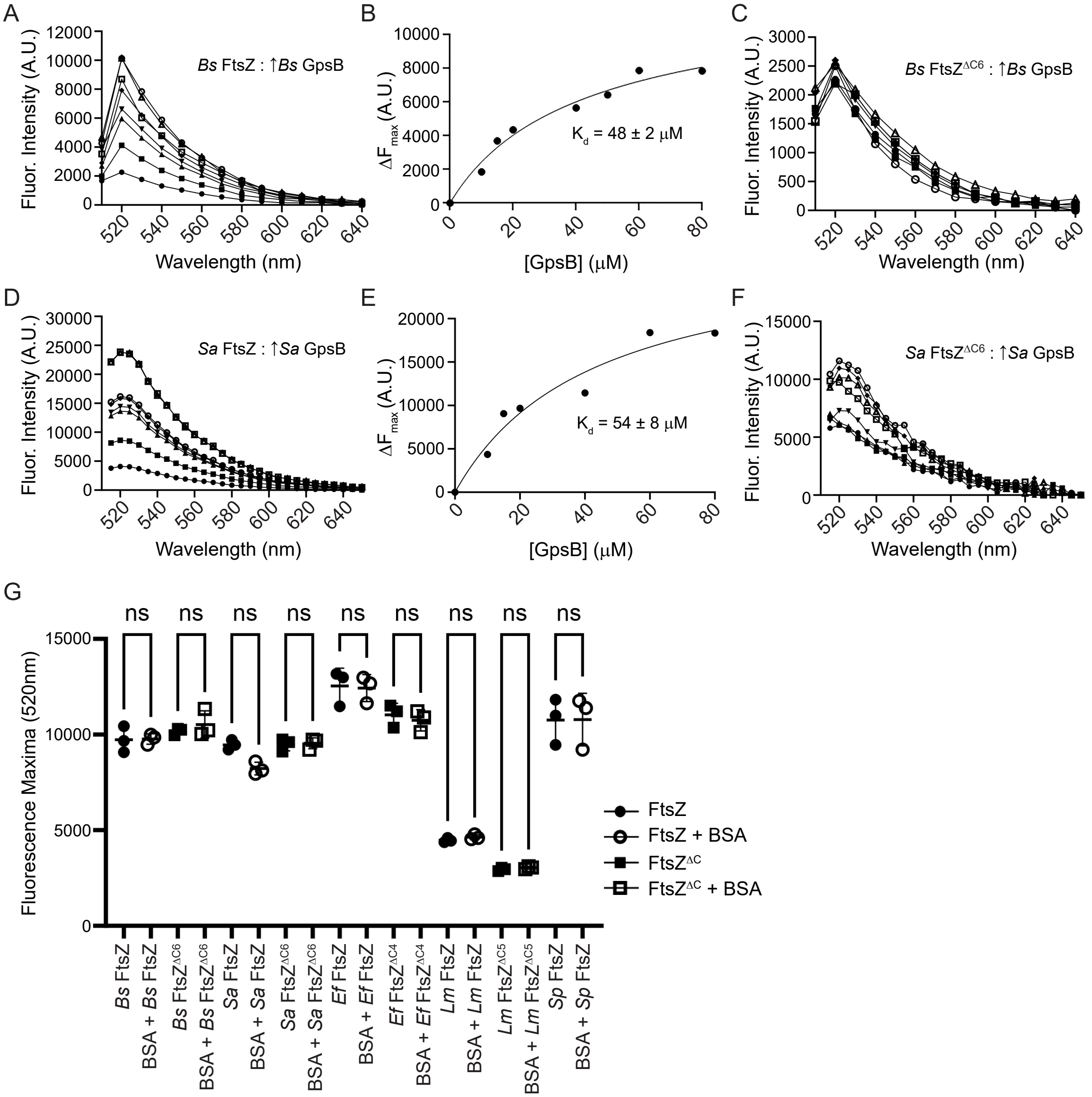
A
Bs FtsZ : ↑Bs GpsB
Fluor. Intensity (A.U.)
Wavelength (nm)
B
K_d = 48 ± 2 μM
ΔF_max (A.U.)
[GpsB] (μM)
C
Bs FtsZ^ΔC6 : ↑Bs GpsB
Fluor. Intensity (A.U.)
Wavelength (nm)
D
Sa FtsZ : ↑Sa GpsB
Fluor. Intensity (A.U.)
Wavelength (nm)
E
K_d = 54 ± 8 μM
ΔF_max (A.U.)
[GpsB] (μM)
F
Sa FtsZ^ΔC6 : ↑Sa GpsB
Fluor. Intensity (A.U.)
Wavelength (nm)
G
ns
Fluorescence Maxima (520nm)
FtsZ
FtsZ + BSA
FtsZ^ΔC
FtsZ^ΔC + BSA
Bs FtsZ
BSA + Bs FtsZ
Bs FtsZ^ΔC6
BSA + Bs FtsZ^ΔC6
Sa FtsZ
BSA + Sa FtsZ
Sa FtsZ^ΔC6
BSA + Sa FtsZ^ΔC6
Ef FtsZ
BSA + Ef FtsZ
Ef FtsZ^ΔC4
BSA + Ef FtsZ^ΔC4
Lm FtsZ
BSA + Lm FtsZ
Lm FtsZ^ΔC5
BSA + Lm FtsZ^ΔC5
Sp FtsZ
BSA + Sp FtsZ

Figure S2

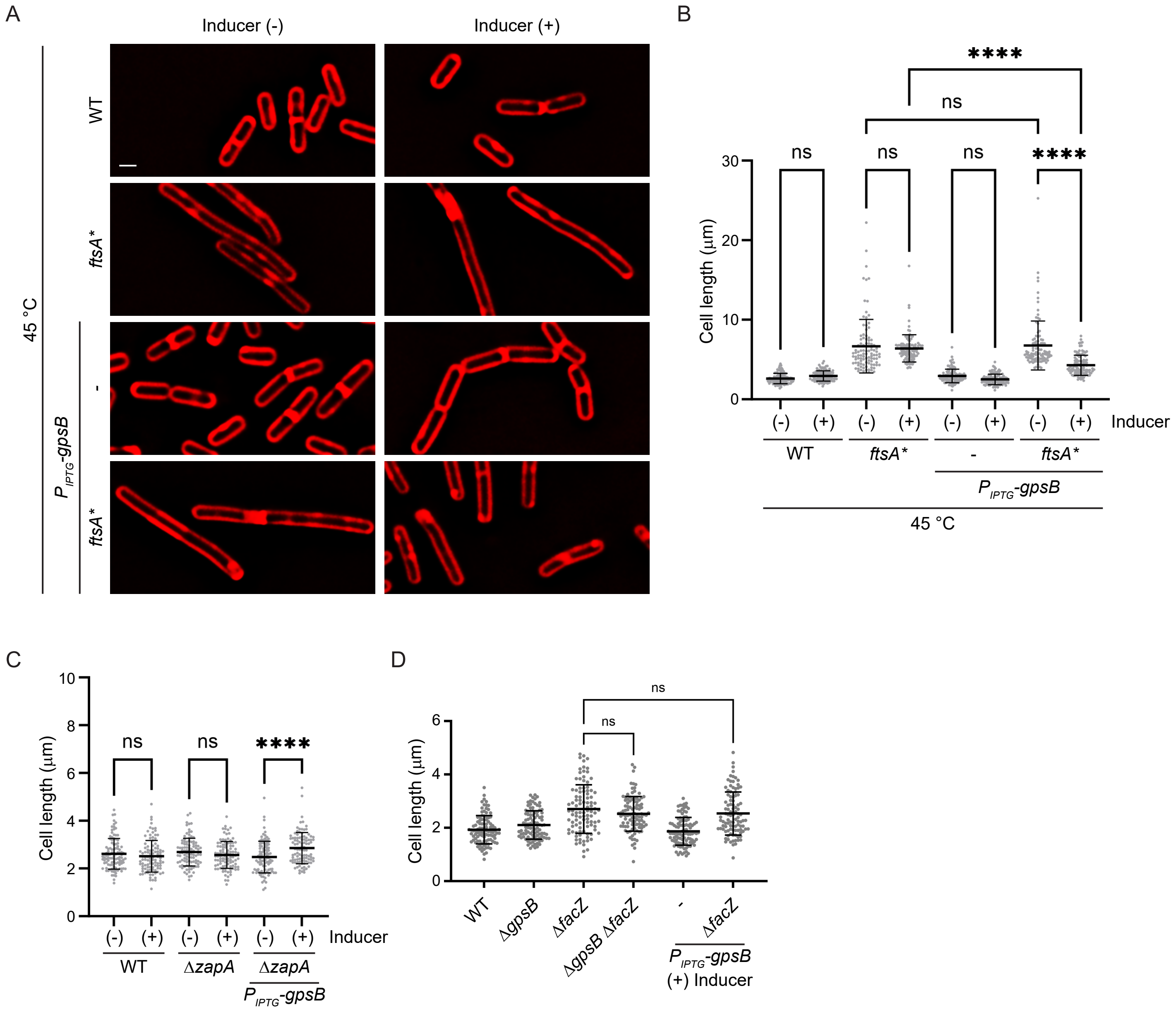

A
Inducer (-)
Inducer (+)
45 °C
WT
ftsA*
$P_{IPTG}$-gpsB
-
ftsA*
B
****
ns
ns
ns
ns
****
Cell length (µm)
30
20
10
0
(-) (+) (-) (+) (-) (+) (-) (+) Inducer
WT
ftsA*
-
ftsA*
$P_{IPTG}$-gpsB
45 °C
C
Cell length (µm)
10
8
6
4
2
0
ns
ns
****
(-) (+) (-) (+) (-) (+) Inducer
WT
ΔzapA
ΔzapA
$P_{IPTG}$-gpsB
D
Cell length (µm)
6
4
2
0
ns
ns
WT
ΔgpsB
ΔfacZ
ΔgpsB ΔfacZ
-
ΔfacZ
$P_{IPTG}$-gpsB
(+) Inducer


Figure S3

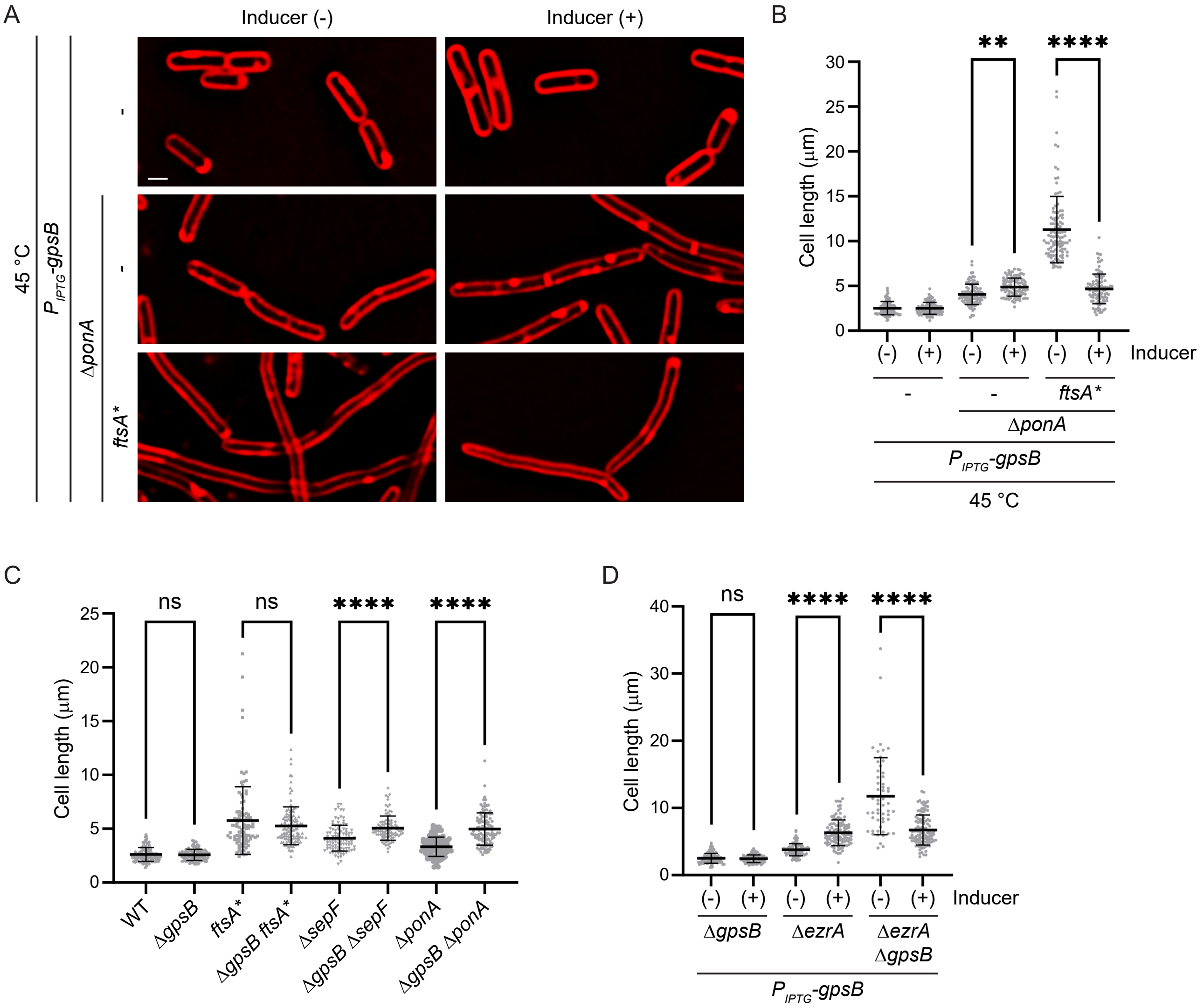
A
Inducer (-)
Inducer (+)
45 °C
P IPTG-gpsB
ΔponA
ftsA*
B
Cell length (μm)
Inducer
ΔponA
P IPTG-gpsB
45 °C
C
Cell length (μm)
ns
****
WT
ΔgpsB
ftsA*
ΔgpsB ftsA*
ΔsepF
ΔgpsB ΔsepF
ΔponA
ΔgpsB ΔponA
D
Cell length (μm)
ns
****
Inducer
ΔgpsB
ΔezrA
ΔezrA ΔgpsB
P IPTG-gpsB

Figure S4
